# Detection of multiple per- and polyfluoroalkyl substances (PFAS) using a biological brain-based gas sensor

**DOI:** 10.1101/2025.09.17.676814

**Authors:** Summer B. McLane-Svoboda, Shruti Joshi, Autumn K. McLane-Svoboda, Camron Stout, Maksim Bazhenov, Debajit Saha

**Author notes:** Corresponding author. (D. Saha).

## Abstract

Per- and polyfluoroalkyl substances (PFAS) are man-made compounds that bioaccumulate in environments. Current PFAS detection technologies encounter difficulty in detecting trace concentrations and require complex data processing, limiting their on-site applicability. By leveraging biological chemical sensing systems (insect olfaction) we can detect broad ranges of PFAS. Insects’ advanced combinatorial coding mechanism at the level of olfactory sensory neurons enables highly sensitive and specific odor detection. Here, we harness the locust olfactory system to differentiate several PFAS. *In-vivo* extracellular neural recordings displayed unique odor-evoked responses for multiple PFAS at environmental concentrations. Using population neuronal response, we classified multiple PFAS with an average 87% accuracy. Machine learning algorithms incorporated separate training and testing datasets, reaching a 61% accuracy. Overall, our study demonstrates the first biological olfaction based broad PFAS detection system.

**Structured Abstract:** *Introduction:* Per- and polyfluoroalkyl substances (PFAS) pose a significant environmental threat due to their widespread presence in consumer waste and resistance to degradation. These “forever chemicals” persist in various ecosystems and exhibit bio accumulative behavior. Increased human exposure to PFAS has been linked to numerous health issues. Despite their growing relevance, current detection methods often lack the sensitivity and efficiency needed for comprehensive environmental monitoring and are unable to simultaneously detect multiple PFAS at environmental concentrations.

*Rationale:* The locust (*Schistocerca americana*) possesses a highly developed olfactory system that has been extensively studied and is accessible for physiological recordings across multiple brain regions. Utilizing olfactory receptor neurons and combinatorial coding strategies, locusts can generate distinct neural fingerprints for trillions of odorants across a wide range of concentrations. Through spatiotemporal neural coding at the antennal lobe (AL) neurons, they can detect chemicals at parts per trillion levels, functioning as a biological chemical sensor with exceptional sensitivity and broad specificity. In this study, we aimed to directly harness the locust’s olfactory neural circuitry to develop a next-generation, brain-based cyborg sensor capable of (1) simultaneously identifying multiple PFAS and (2) determining their concentration ranges, addressing key limitations of existing detection technologies with the integration of machine learning algorithms.

*Results:* In-vivo extracellular neural recordings from the locust AL revealed that individual neurons exhibited distinct response profiles to different PFAS and their varying concentrations, indicating that neuronal activity if modulated by both chemical identity and concentration. By incorporating both spatial and temporal dimensions of neural activity, the recorded neuronal populations produced unique and reproducible response patterns corresponding to specific PFAS and concentration levels. Using this approach, the cyborg sensor demonstrated an overall detection accuracy of 87% across a panel of seven PFAS, with high sensitivity and specificity for both individual analytes and broader chemical groupings. Notably, classification of PFAS concentration ranges down to parts per trillion achieved 84% accuracy, with PFOS concentrations reaching 100% detection rate. A machine learning algorithm trained on high concentration data and tested on low concentration data achieved a 61% accuracy. These results underscore the potential of biologically integrated cyborg sensors for real-time, high-resolution environmental monitoring of several PFAS.

*Conclusion:* This study demonstrates, for the first time, the locust olfactory neural network harnessed as a highly effective cyborg sensor for detecting and classifying various PFAS across concentration ranges. This sensor can accurately distinguish between multiple PFAS and controls with high sensitivity and specificity. Through combinatorial coding and spatiotemporal neural dynamics, the locust neural computation encodes distinct activity patterns in response to PFAS and their concentrations. These neural signatures serve as unique “fingerprints” for individual PFAS and concentrations, enabling precise identification. *In-vivo* electrophysiological recordings revealed clear, compound-specific differences in neural activity, with high classification accuracy. Real-time and machine learning analysis further addressed key limitations of conventional PFAS sensors. This novel approach represents a significant step toward the development of compact, real-time, brain-based PFAS detection sensor capable of discriminating multiple compounds and concentrations simultaneously.

## Main Text

Per- and polyfluoroalkyl substances (PFAS) are chemically persistent compounds that are found ubiquitously in the global environment, each containing at least one carbon atom that is fully fluorinated(*1–3*). Often referred to as “forever chemicals” due to their durability and resistance to degradation, PFAS are present in a variety of household items, contributing to increased human exposure(*4, 5*). PFAS pose significant health risks due to their bioaccumulation in environments(*1, 6–8*) animals(*9*), agricultural crops(*10*), and drinking water(*11*). While a comprehensive record of the adverse health effects from PFAS consumption is still unknown(*12, 13*), research suggests that the accumulation of these compounds are linked to a range of diseases(*14–20*) and immunotoxicity(*21*), including potential carcinogenic effects(*22*). This creates an urgent need for more advanced chemical sensors capable of precise environmental pollution detection.

Conventional gas sensors, such as liquid and gas chromatography-mass spectrometry (LC-MS and GC-MS), encounter difficulty with detecting chemicals at ppt levels, require extensive data processing, and lack portability; making them impractical for fieldwork(*23*). While hybrid systems like electronic noses and molecularly imprinted polymers (MIPs) attempt to replicate biological olfaction using pattern recognition algorithms, are limited in sensitivity, typically detecting a few compounds at ppb levels and focusing on single-compound detection(*23, 24*). Conversely, biological gas sensing systems (i.e., locust olfaction), can detect chemicals at environmental concentrations by employing a highly evolved advanced combinatorial coding scheme to differentiate between odorants and concentrations, making them the ideal candidate for PFAS detection and identification(*24, 25*). Locusts possess a well-developed olfactory system that is accessible for physiological recordings at multiple stages of neural processing(*26*). Their antennae house numerous olfactory receptor neurons (ORNs), which respond to diverse odorant stimuli, triggering a transformation of stimuli into odor-evoked electrical signals (neural spikes)(*27, 28*). These signals are transmitted to the antennal lobes (ALs), where excitatory projection neurons (PNs) and inhibitory local neurons process the information before passing it to higher order brain centers(*27*). The AL network generates unique neural fingerprints for odorants and concentrations using a spatiotemporal coding scheme, enabling a precise gas sensor capable of identifying trillions of odorants at ppm - ppt levels(*29*), without the need for training and binary behavioral responses (e.g., canine olfaction)(*23*). Previous studies have shown that the locust is highly effective in distinguishing various biological compounds (i.e., oral cancers, associated biomarkers, endometriosis, and bacterial biofilms)(*27, 30–32*). Building on this foundation, we conducted *in-vivo* extracellular AL neural recordings using the locust olfactory system to detect and classify multiple PFAS at environmental concentrations. A machine learning (ML) mechanism was utilized to detect and identify PFAS at various concentrations. We hypothesized that the locust olfactory network could accurately identify PFAS simulating environmental concentrations via changes in neural activity. Through this forward-engineering approach, we aim to develop a highly sensitive, detection method specific to PFAS, addressing the limitations of current technology.

### PFAS are detected by the olfactory neurons in the locust AL

We measured odor-evoked neural responses from the locust olfactory circuitry to seven pure PFAS and an odor control (room air). The primary objective of these experiments was to investigate the broad detection range of PFAS. In brief, a multichannel electrode was inserted into one of the locusts ALs; odor delivery and airflow were controlled using a commercial-grade olfactometer (Materials and Methods, **Fig. 1A and S1**). The stimuli were presented in a pseudorandomized order across five trials to avoid temporal biases. *In-vivo* electrophysiological data was recorded and voltage traces were spike-sorted to identify individual neurons that responded to the PFAS odor stimuli (Material and Methods).

**Fig. 1.**
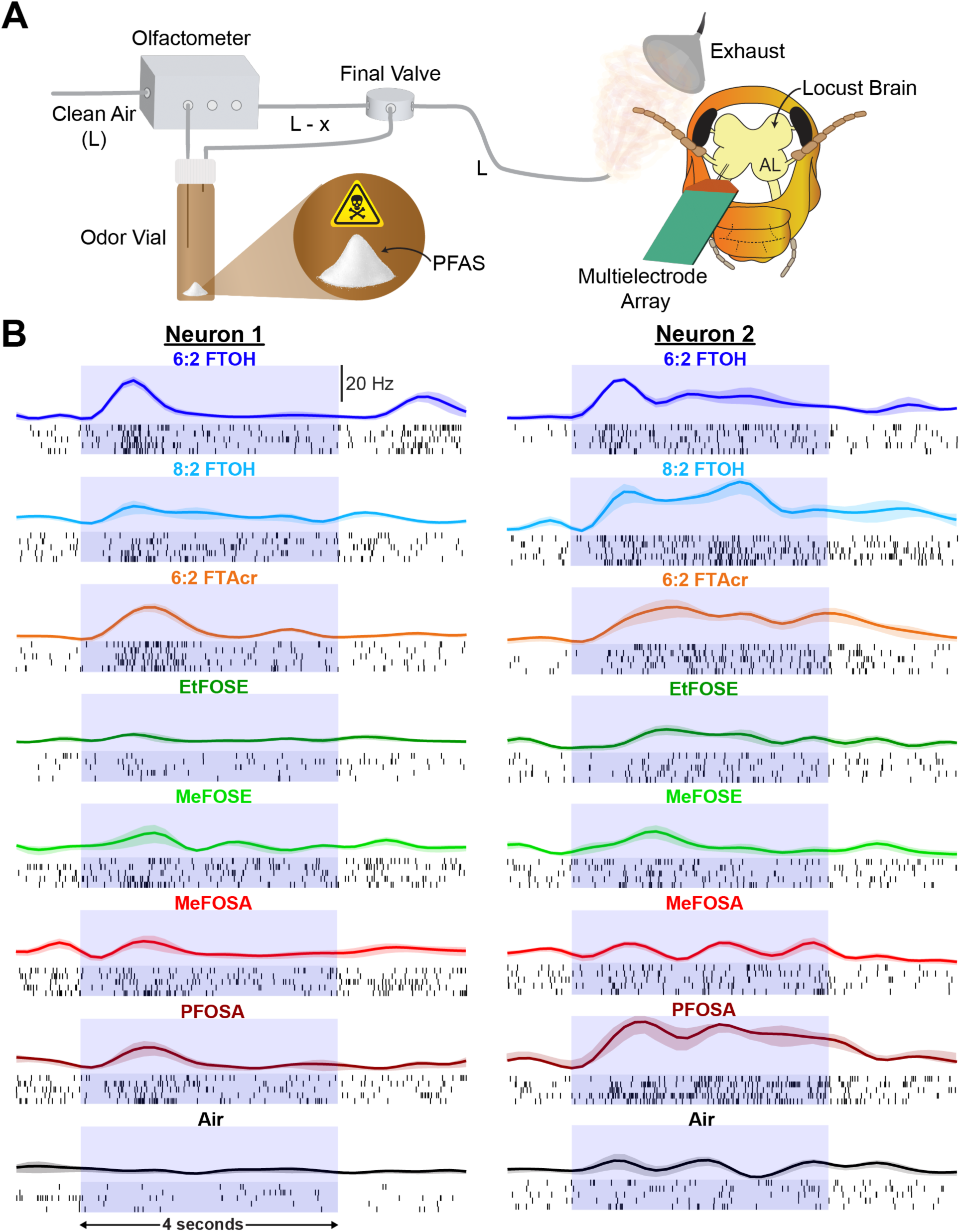
Unique odor-evoked neural responses to pure PFAS. **(A)** A schematic illustrates the odor delivery system used for in-vivo neural recordings. Pure PFAS were placed into airtight odor vials. A fixed volume of clean air (L) from a compressed air cylinder flowed into the olfactometer, passing through the odor vial to collect volatile compounds from a single PFAS. The resulting odor-laden air mixture (L-x) was then delivered to the locust antennae. The experimental setup-controlled odor volume and duration. Neural activity was recorded from the AL using a multichannel multielectrode array. An exhaust system removed any residual odor. **(B)** Extracellular voltage traces were spike-sorted to extract neural responses. Two representative neurons displayed distinct odor-evoked activity patterns for all tested PFAS. Raster plots and peri-stimulus time histograms (PSTHs) were generated for each neuron, showing responses to seven PFAS and an empty control vial. Each black line in the raster plots represents a spiking even across five trials per odor condition. PSTHs generated from the raster plots illustrate changes in firing rate over the stimulus duration, with shaded regions indicating the stand error of the mean (S.E.M.). The light blue box marks the odor stimulus window (4 seconds).

We found that odor-evoked neural activity from a single extracted neuron varies in response to all seven PFAS and the control (**Fig. 1B**). Raster plots depicted individual action potentials across five trials. Averaging the trials allowed us to visualize the firing frequency of one neuron in peri-stimulus time histograms (PSTHs), revealing changes in firing frequency throughout odor stimulation (ON-response), as shown by the light blue boxes (**Fig. 1B**). Two representative neurons demonstrated distinct differences in responses to multiple PFAS and the control (**Fig. 1B**). Each neuron in the population exhibited a unique response to the odors, prompting examination of the neuronal population’s response. We compiled 86 neurons (n = 86) to investigate the population response. Analysis displayed the collective neural response to each stimulus, with distinct ON- and OFF-responses (post-stimulus) observed for certain PFAS (**Fig. 1B**). These findings suggest that the neuronal population responds to PFAS through a combination of ON and OFF responses, creating a unique neural fingerprint for each PFAS. The most variance in the odor-evoked neural data was observed between 0.25 – 1.0 seconds post-stimulus onset, likely due to a delay in odor response following the final valve opening.

### Population neural responses can differentiate between multiple PFAS

We applied dimensional reduction techniques to examine the neuronal populations response between various PFAS (Materials and Methods). Neurons from electrophysiology recordings (n = 86) were combined to generate an odor-evoked population response matrix (*neurons x time*). Responses were aligned at stimulus onset and averaged across five trials. Firing rates for each neuron were binned into non-overlapping, 50 ms time intervals, chosen based on 20 Hz oscillations observed in the insect ALs(*33, 34*). The number of spikes within each time bin was counted to create odor-specific, high-dimensional population response vectors. Principal component analysis (PCA) was used to visualize the temporal odor-evoked responses of the neuronal population. PCA was performed using a 750 ms time window from 0.25 to 1.0 seconds post-stimulus onset to capture the period of highest odor-evoked response. The three principal components with the highest eigenvalues, which explained the most variance in the dataset, were projected onto three dimensions for visualization (**Fig. 2A**). Odor-specific response vectors were connected across adjacent time bins, forming stimulus neural trajectories. Trajectories were aligned at the origin, represented by a black dot. Notably, each odor stimulus produced distinct neural trajectories, indicating unique neuronal population responses to all PFAS, as well as the control (**Fig. 2A**). Further analysis using root mean square (R.M.S.) for real-time data analysis revealed similar results (n = 51, **fig. S2A**). Additionally, we applied a second dimensionality reduction technique – supervised linear discriminant analysis (LDA) – to maximize variance between stimulus clusters while minimizing variance within each stimulus. We observed clear neuronal clustering for each PFAS in neuronal populations and in R.M.S. analyses (**Fig. 2B, fig. S2B**). These findings suggest that the spatiotemporal AL neural responses can discriminate across a range of PFAS.

**Fig. 2.**
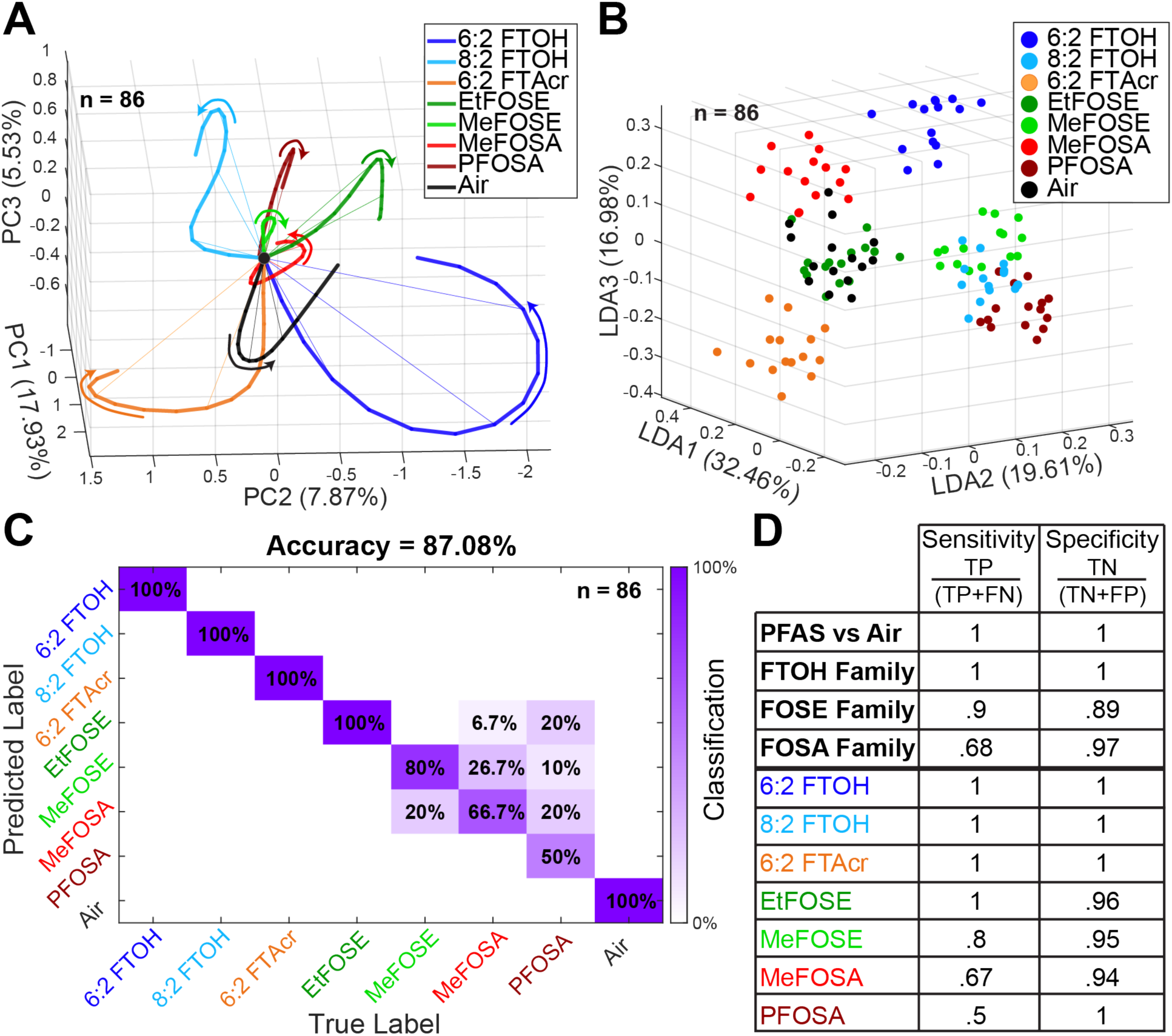
Classification of pure PFAS using population neural responses. **(A)** PCA was used to reduce dimensionality and visualize population PN responses to seven pure PFAS and a control. Trajectories represent odor-evoked neural activity between 0.25 – 1.0 seconds after odor exposure. Colored arrows indicate the evolution of these trajectories, with all trajectories aligned at 0.25 seconds after odor onset (black dot), which corresponds to the delay in odor delivery after opening the final valve (0.0s). A greater angular trajectory from the origin and the other odors indicates a more distinct neural response. A total of 86 neurons were analyzed. **(B)** LDA, a supervised dimensionality reduction technique, was applied to classify odor responses between 0.25 – 1.0 seconds post-exposure. LDA separates responses into eight distinct clusters, minimizing within-cluster variance while maximizing separation between different odors. This clustering demonstrates how PFAS influence neural temporal dynamics over the analyzed timeframe. **(C)** A high-dimensional LOTO confusion matrix was used to classify responses to seven PFAS and the control vial (Materials and Methods). The matrix compares true versus predicted labels, with the darkened diagonal indicated correct classifications based on the smallest Euclidean distance. Using the entire response window (0.25 – 5.75 seconds), which includes both odor onset and offset responses, the classification achieved an accuracy of 87.08%. This result confirms that our sensor can reliably detect and distinguish between different PFAS. **(D)** Sensitivity and specificity table highlighting precise compound detection using the locust sensor. Values were calculated from the LOTO matrix (Materials and Methods).

Quantitative classification was performed using a leave-one-trial-out (LOTO) matrix (Materials and Methods). In brief, the neuron response was aligned and divided into 50 ms non-overlapping time bins. The number of spikes within each time bin was counted to create a 3D matrix (*trials x neurons x time bins*). Four out of five trials were combined to form a training template, while the remaining trial formed a testing template. The templates were plotted in high-dimensional space; each testing template was assigned to the closest training template based on Euclidean distance. This procedure was repeated for each trial serving as the testing template once. The classification results were determined by the mode of time bin assignments, with higher classification accuracy observed along the diagonal of the matrix, where testing templates were correctly matched to the corresponding training templates. A time window from 0.25 to 5.75 seconds post-stimulus onset (5,500 ms) was used to capture both ON and OFF responses. The classification success rate for the neuronal population (n = 86) was 87.08% (**Fig. 2C**). Notably, five out of eight PFAS achieved 100% classification accuracy, while success rates for other stimuli were lower (**Fig. 2C**). R.M.S. data (n = 51) yielded an 85% classification success rate, with 100% accuracy for four of the PFAS (**fig. S2C**).

To further assess the precision of PFAS classification, sensitivity and specificity values were calculated (Materials and Methods). These values were derived from the LOTO confusion matrix, where columns represented true positives (TP) and false negatives (FN), and rows represented false positives (FP) and TP. Any data outside the relevant odor column or row was considered a true negative (TN). True positives were identified as the diagonal classification accuracy, where the true label was correctly assigned to the predicted label. We observed that most PFAS achieved sensitivity and specificity values of 90% or higher, with three odors exhibiting lower sensitivity values (**Fig. 2D**). Additionally, when discriminating between PFAS families (e.g., FTOH, FOSE, FOSA), all families achieved sensitivity and specificity values of approximately 90% or higher, except for one, which showed reduced sensitivity (**Fig. 2D**). When distinguishing between PFAS and air (non-PFAS), the sensor demonstrated 100% precision (**Fig. 2D**). For real-time R.M.S. analysis, a 90% sensitivity and specificity were obtained for all but three PFAS and the FOSA family, showing decreased sensitivity (f**ig. S2D**). Our novel cyborg gas sensor can accurately differentiate between multiple PFAS, PFAS families, and controls.

### Population neural responses can differentiate environmental PFAS concentrations

We examined neural responses to a subset of PFAS at environmental concentrations (i.e., ppb and ppt) to investigate the sensors detection limit. For the PFAS tested – 6:2 FTOH and 6:2 FTAcr, both receiving 100% classification in previous experiments – two concentrations were selected that mimic environmental conditions (0.01% vol/vol and 0.0001% vol/vol diluted in ddH2O). The previously high classification accuracy, along with strong odor-evoked responses, made these compounds ideal for examining neural responses at environmental levels. PPB concentrations were calculated using Raoult’s Law, with ppt levels observed at the lowest concentration (**Fig. 3A**) (Materials and Methods). Odor stimuli were delivered to the locust antennae using previously described methods (**fig. S1**, Materials and Methods).

**Fig. 3.**
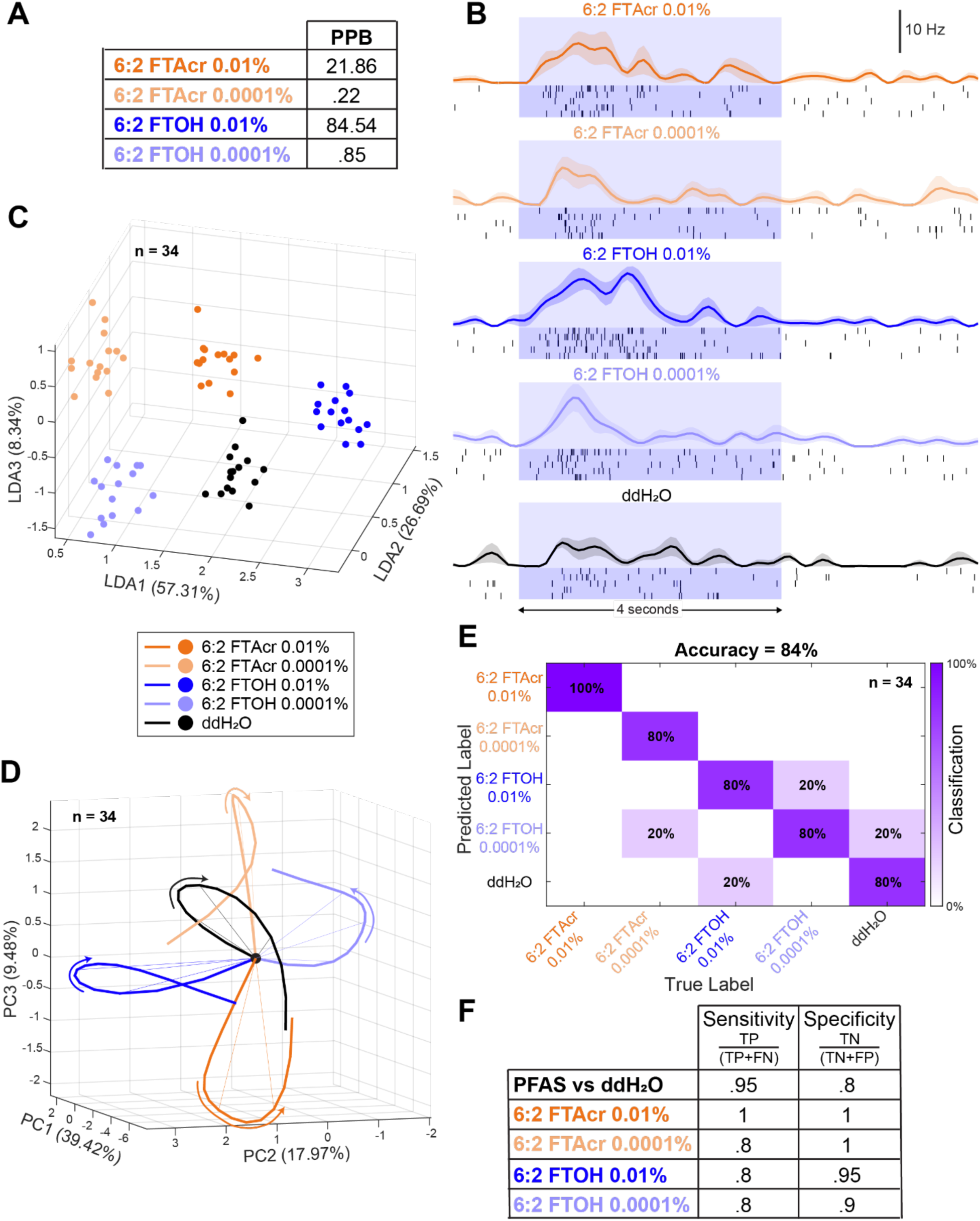
Real-time classification of environmental PFAS concentrations using R.M.S. **(A)** Raoult’s law was used to calculate the ppb concentration ranges for PFAS. The lowest concentration (0.0001%) represents ppt ranges, which mimic those typically found in environmental samples. **(B)** VOC-evoked extracellular neural voltage traces for representative neurons exposed to PFAS concentrations were spike-sorted. Spike timing is shown in raster plots across five trials, with each black line indicating a single spiking event. PSTHs generated from the raster plots reveal changes in spiking patterns, with shaded regions representing the SEM. Light blue boxes mark the odor stimulus windows (4 seconds). **(C)** A 3-class LDA was used to separate R.M.S. odor responses between 0.25 – 1.0 seconds post-stimulus onset. Five distinct clusters were visualized, demonstrating clear spatiotemporal separation between different odors and concentrations. A total of 34 tetrodes were used in the analysis. **(D)** R.M.S. neural trajectories of two PFAS at two concentrations, along with a control odor (5 odors total), were visualized using PCA dimensionality reduction. The VOC-evoked responses are plotted between 0.25 – 1.0 seconds after stimulus onset. Colored arrows indicate the trajectory evolution of responses. Trajectories are aligned at 0.25 seconds after odor onset (indicated by a black dot), corresponding to the delay after the final valve is opened. An increased angular trajectory from the origin and other odors signifies a distinct neural response. A total of 34 tetrodes were analyzed. **(E)** A classification matrix based on the LOTO analysis shows 84% classification accuracy within the 0.25 – 5.75 seconds time window after stimulus onset, demonstrating that the sensor can effectively detect PFAS concentrations. **(F)** A sensitivity and specificity table highlights the precise detection of PFAS and their concentrations. Values were calculated from the LOTO matrix (Materials and Methods), demonstrating the sensor’s ability to accurately classify PFAS odors and their concentrations.

Interestingly, recordings from multiple ALs revealed that neurons responded differently to PFAS concentration and the odor control. A representative neuron demonstrated stimulus-specific changes in spiking activity, visualized via raster plots and corresponding firing frequency shifts shown in PSTHs (**Fig. 3B**). This data suggests that the sensor can detect PFAS odors at trace concentrations. To further characterize neural dynamics, R.M.S. analysis was applied using discrete 50 ms time bins to visualize and classify responses in real-time (n = 34). The most significant neural activity changes occurred between 0.25 – 1.0 seconds following stimulus onset. LDA revealed clear separability of neural response clusters for each PFAS, its concentrations, and the control, indicating distinct spatiotemporal encoding for each condition (**Fig. 3C**). PCA further demonstrated unique neural trajectory evolutions over the course of odor presentation (n =34, **Fig. 3D**). Odor-specific response vectors, linked across consecutive 50 ms bins, formed distinct trajectories, all aligned to a common origin. The divergence of these trajectories from the origin and other stimuli underscores the specificity of the voltage trace responses. Overall, these results indicate that the cyborg sensor can detect and encode differences in concentration of the same PFAS, enabling discrimination through spatiotemporal neural activity patterns.

Classification success rates remained high for the both PFAS across varying concentrations during quantitative analysis. A high-dimensional LOTO analysis was employed to assess odor classification accuracy (Materials and Methods). A post-stimulus time window of 0.25 – 5.75 seconds was used for trial-wise classification, encompassing both ON and OFF phases of neural population responses (n = 34). Within this window, classification accuracy reached 84% (**Fig. 3E**). The sensor consistently classified all PFAS and concentrations with an accuracy of 80% or higher (**Fig. 3E**). Additionally, sensitivity and specificity analyses displayed values above 80% for all stimuli, including the comparison of PFAS versus control. A slight drop in sensitivity was observed for the lowest concentration (0.0001%) (**Fig. 3F**). These findings quantitatively demonstrate that the cyborg sensor can reliably distinguish between different PFAS and their concentrations at environmental levels (i.e., ppb and ppt ranges).

### Population responses can differentiate PFOS at environmental concentrations

Stimuli tested thus far have primarily included volatile PFAS with measurable vapor pressures. While these compounds are detectable by the locust olfactory system at environmental concentrations, they are not the most commonly found PFAS(*35–37*). Perfluorooctanesulfonic acid (PFOS), one of the most environmentally prevalent PFAS, is known for its extremely low vapor pressure, making it effectively non-volatile and challenging to detect via gas sensing(*38*). To investigate whether the brain-based sensor can detect common but very low-volatile PFAS, we tested PFOS at environmental concentrations (0.01% v/v and 0.0001% v/v). Ppb and ppt values were calculated using Raoult’s law and known physicochemical properties of PFOS (Materials and Methods). The lowest concentration tested (0.0001% v/v) represented approximately 34 ppt, simulating real-world environmental conditions (**Fig. 4A**). Compared to previously tested PFAS, PFOS concentrations represent lower vapor-phase levels (**fig. S3A**). Odor stimuli were delivered pseudo randomly to the locust antennae and odor-evoked responses were recorded from multiple ALs. Distinct differences in neural activity were observed between PFOS concentrations and the odor control. A representative neuron demonstrated concentration-specific spiking patterns in raster plots and corresponding PSTHs (**Fig. 4B**). Neuronal responses were distinguishable from the odor control and between PFOS concentrations, indicating that the locust neural circuitry can discriminate minute PFOS concentrations, independent of the background matrix (**Fig. 4B**).

**Fig. 4.**
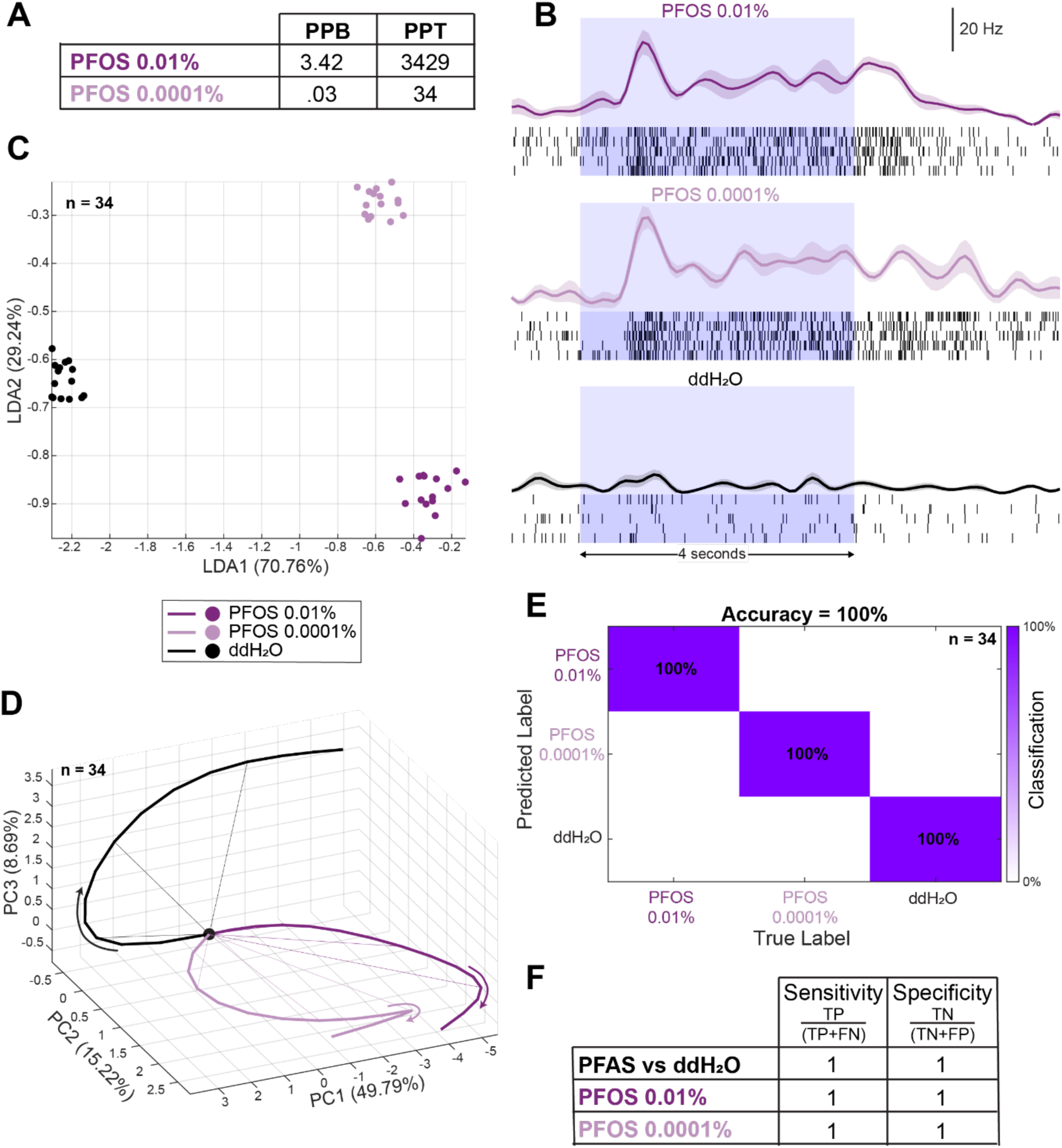
Precision classification of PFOS at environmental concentrations using R.M.S. **(A)** PFOS concentrations in ppb and ppt were calculated using Raoult’s law, based on chemical characteristics found in the literature (Materials and Methods). The lowest concentration represents ppt values. **(B)** Extracellular voltage traces were spike-sorted to reveal individual neurons. Raster plots show spike timing across five trials, where each black line represents a spiking event. The average firing frequency across the five trials is displayed in the PSTHs, with shaded regions representing the SEM. Blue boxes indicate the odor stimulus windows (4 seconds). PSTHs illustrate variations in odor-evoked responses across different concentrations and the control. **(C)** A 2-class LDA dimensionality reduction technique was applied to visualize R.M.S. neural response clustering between 0.25 – 1.0 seconds after odor presentation. A total of 34 tetrodes were used in the analysis. **(D)** PCA dimensionality reduction analysis highlights distinct odor trajectory evolution for two PFOS concentrations, along with a control, between 0.25 – 1.0 seconds after stimulus onset. The separation between PFOS and the control, as well as differentiation in neural response trajectories between concentrations, shows that the locust can distinguish between low PFOS concentrations. Trajectories are aligned at 0.25 seconds after odor onset (black dot), with colored arrows indicating trajectory evolution. Distinct neural responses are shown by increased angular deviation from the origin and separation from other odors. A total of 34 tetrodes were analyzed. **(E)** A LOTO classification matrix demonstrates 100% classification accuracy within the 0.25 – 5.75 seconds time window after stimulus onset. **(F)** A sensitivity and specificity table shows 100% precision in detecting environmental PFOS concentrations. Values were calculated from the LOTO matrix (Materials and Methods).

To visualize odor-evoked neural responses in real-time, we applied a R.M.S. analysis using non-overlapping 50 ms time bins (n = 34). The most significant changes in neural activity occurred between 0.25 – 1.0 seconds after stimulus onset. Using LDA, we observed clear separation of neural clustering into three distinct groups (**Fig. 4C**). Notably, the odor control formed a cluster in a spatial region opposite from those representing PFOS responses, indicating that the locust olfactory system uses different neural spatial codes to represent concentrations of PFOS. When combining PFOS data with previously tested PFAS concentrations, high-concentration PFAS (0.01%) and the odor control each formed well-separated clusters, while lower concentrations (0.0001%) showed more spatial overlap (**fig. S3B**). Despite this overlap, the 0.0001% responses still exhibited partial separation, suggesting that the sensor can differentiate multiple PFAS at ppt levels. PCA revealed distinct neural trajectory evolutions for the control versus PFOS stimuli (**Fig. 4D**). The control trajectory diverged separate from PFOS concentrations. Both PFOS concentrations showed greater expansion from the origin, reflecting active and differentiated neural responses to low concentrations. Additionally, differences in trajectory evolution were evident between the PFOS concentrations (**Fig. 4D**). When integrating PFOS trajectories with previous PFAS, each PFAS exhibited distinct trajectories (**fig. S3C**). Controls consistently followed a trajectory opposite to PFAS. While trajectories of the same compound at different concentrations were generally similar, they retained distinct differences. These findings emphasize that the locust neural circuitry encodes unique signatures for environmental PFOS concentrations, distinguishable from other PFAS and the control.

For quantitative classification, a high dimension LOTO analysis was used. Trial-wise classification was performed using a 0.25 – 5.75 second post-stimulus window, encompassing both ON and OFF phases of neural activity (n = 34). Analysis achieved a 100% classification success rate, accurately distinguishing environmental PFOS concentrations and the control (**Fig. 4E**). Sensitivity and specificity values showcased PFOS concentrations and PFAS versus control comparison achieving 100% (**Fig. 4F**). When combining PFOS data with previous PFAS responses, the overall classification accuracy remained high at 85.71% (n = 34, **fig. S3D**). All PFAS concentrations achieved an accuracy of 80% or higher, with the exception of 0.0001% 6:2 FTOH. Sensitivity and specificity for the combined dataset was greater than 80% across PFAS concentrations, with minor decreases in 0.0001% 6:2 FTOH sensitivity (**fig. S3E**). Findings indicate that the sensor can reliably distinguish between PFOS and other PFAS at environmental concentrations. The brain-based cyborg sensor exhibits high sensitivity and specificity, making it a powerful tool for detecting trace levels of PFOS and PFAS.

### Machine learning algorithm utilized for identification and cross-concentration generalization of PFAS

When classifying odor identity independent of concentration, the CNN-LSTM hybrid model achieved a performance of 78.5% accuracy with a standard deviation of 0.04% across different random initializations (**Fig. 5A**). This reflects the model’s ability to classify compounds across odor concentrations as long as the training templates contained a subset of low concentration data. The per-class performance analysis revealed that 6:2 FTAcr achieved 83% accuracy and 6:2 FTOH showed 86% accuracy across concentrations, while PFOS demonstrated 95% accuracy for both concentration levels tested. Water controls presented a challenge with 50% accuracy.

**Fig. 5.**
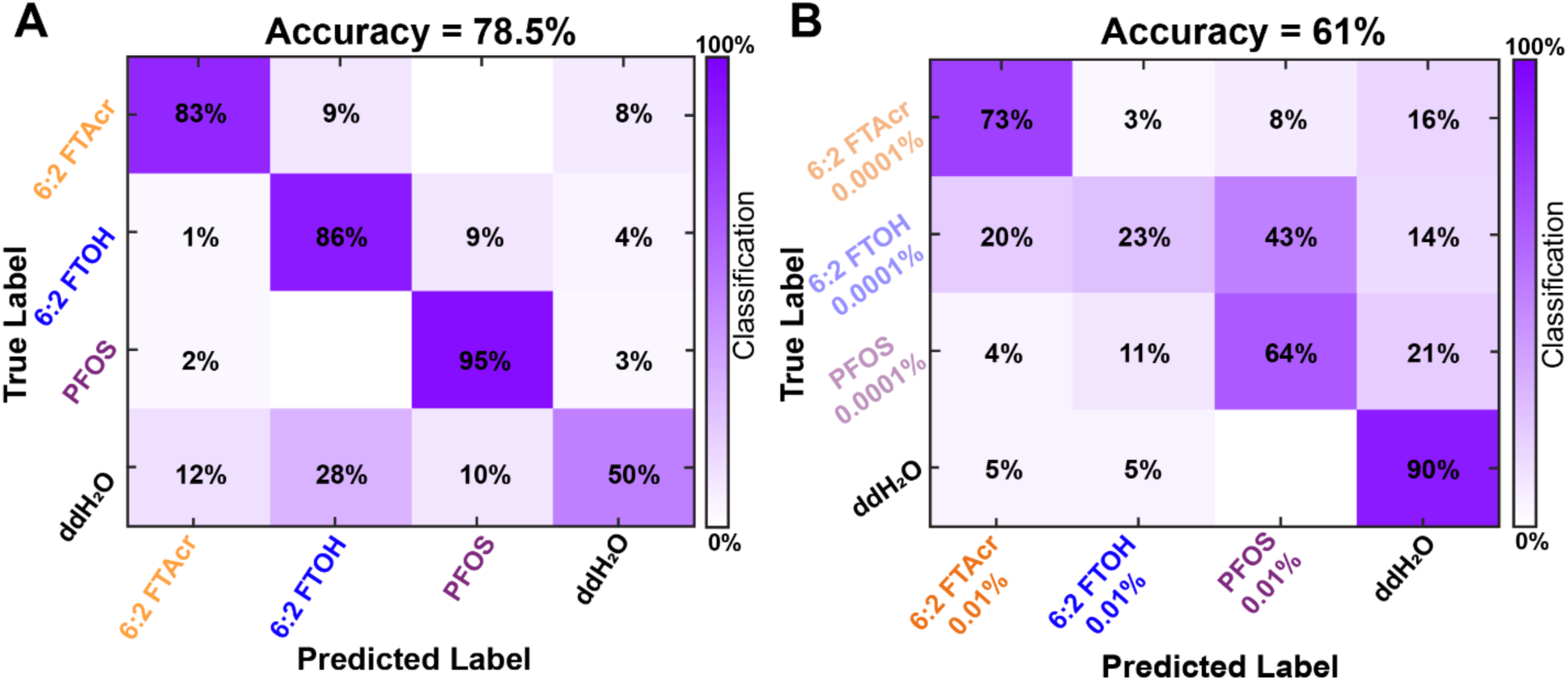
Performance of machine learning model. **(A)** Confusion matrix for classification of PFAS independent of concentration. Average classification accuracy reached 78.5% for the 0.25 – 5.75 second time window. **(B)** Classification confusion matrix for different compounds with hybrid CNN-LSTM model trained on neural responses to high concentration of the compound and tested on low concentrations of the compound. The matrix demonstrates a 61% accuracy within the 0.25 – 5.75 second time window after odor stimulus.

The CNN-LSTM hybrid model was able to achieve cross-concentration generalization, containing a 61% average accuracy with a standard deviation of 0.2% across random initializations when trained exclusively on high-concentration data and tested on low-concentration samples (**Fig. 5B**). The per-class generalization performance revealed significant differences across the different compounds, with 6:2 FTAcr achieving 73% accuracy, PFOS achieving 64% accuracy, and 6:2 FTOH showing 23% accuracy. Water was represented by an identical neural response in both training (high concentration) and testing (low concentration) sets, leading to a correspondingly high classification accuracy. The hybrid CNN-LSTM model’s success stems from its ability to extract local spatiotemporal relationships through the convolutional component and global spatiotemporal dynamics through the recurrent component, effectively capturing concentration-invariant features of odor identity.

## Discussion

The detection of PFAS has gained increasing importance in recent years. These substances are found across diverse natural environments, leading to growing human exposure and bioaccumulation in both the environment and living organisms(*39–41*). As a result, PFAS exposure has been linked to a range of health conditions(*16, 20–22, 42*). This study focuses on leveraging the neural computations of biological olfaction to detect multiple PFAS at environmental concentrations, varying depending on the extent of environmental pollution. We demonstrate distinct neural activity patterns in response to individual PFAS, as well as variations in neural responses to different concentrations of the same compound. This is the first study to utilize a brain-based, cyborg sensor to differentiate PFAS identity and concentration. Current detection technologies lack the combination of broad sensitivity and enhanced specificity required for this level of discrimination.

Traditional gas sensors (i.e., LC-MS and GC-MS) lack the ability to detect ppt concentrations due to the need for extensive data processing, making them unsuitable for field applications(*23*). Most GC-MS systems detect total fluorine content without identifying PFAS or specific concentrations(*43, 44*). Hybrid systems (e.g., e-noses and MIPs) aiming to imitate biological olfaction detect only a few compounds at high concentrations simultaneously(*23, 24*). Identifying multiple PFAS and their concentrations would require several calibrated MIPs, as many PFAS lack established standards(*45*). Biological detection systems address these limitations by offering high sensitivity and broad specificity. While biology has previously been applied to PFAS detection (e.g., using bacterial cultures to analyze gene expression in response to contamination) these methods rely on genetically engineered organisms and involve significant post-processing(*46, 47*). In contrast, little research has explored biological olfaction for PFAS detection. Honeybee olfaction, shown to neurally distinguished lung cancers(*24*), has been studied in the context of PFAS avoidance, relying on behavioral assays and extensive training that yields binary outputs without identifying specific chemicals presented(*48*). Analyzing the underlying biological neural mechanisms offers a promising path toward chemical detection without the need for behavioral training. By integrating the precision of biological olfactory neural circuits with modern classification algorithms, our cyborg sensor merges the accuracy of natural systems with the scalability and practicality of engineered technologies.

Previous work has demonstrated the locust olfactory system accurately distinguishing between a variety of cancers, endometriosis, and man-made substances (e.g., DNT and TNT) through AL neural recordings(*25, 27, 28*). In this study, we extend this approach to detect different PFAS and their concentrations. By leveraging the locust olfactory network, we utilize it as a novel part-brain cyborg sensor to accurately classify multiple PFAS, addressing limitations of existing devices and marking a significant advancement in PFAS detection. The primary advantage of this system lies in biological olfaction’s ability to generalize diverse chemicals, arising from the advanced spatio-temporal coding system where time and space contribute to stimuli neural dynamics. Our findings showcase a highly sensitive and specific biological cyborg sensor capable of distinguishing multiple PFAS at environmental concentrations. Individual neurons respond to minute changes in PFAS identify and concentrations, compared to control odors (**Figs. 1-4**). We observed differences in neural spiking frequency across varying concentrations of PFAS as well (**Figs. 3 and 4**). These variations correspond to unique spatio-temporal responses and distinct encoding patterns for PFAS identity versus concentration, highlighting the sensor’s precise ability in detecting multiple PFAS. Furthermore, PFOS is the most abundant PFAS found in the environment, characterized by its non-volatile properties and presence at ppt concentrations(*49, 50*). In this study, we present a novel method for detecting PFOS at environmental concentrations in real-time, revealing neural activity changes to PFOS relative to baseline, with clear distinctions across concentrations (**Fig. 4B**). We identified spatio-temporal encoding patterns that accurately distinguished environmental PFOS concentrations in real-time (**Figs. 4C-F**). This is the first study to demonstrate a brain-based cyborg gas sensor capable of detecting a broad range of PFAS, including PFOS, at ppt levels in real-time. We developed a ML algorithm for cross-concentration generalization (**Fig. 5**). This algorithm uses high-concentration data to successfully identify compounds at low concentrations. The ML approach displays generalization across PFAS concentrations, identifying compounds at environmental concentrations. This approach addresses a critical challenge in environmental monitoring where the training data may not encompass the full range of concentrations encountered in real world applications. The varying generalization performance across different PFAS provides insights into biological odor processing mechanisms. PFOS and 6:2 FTAcr show robust concentration-invariant representation, suggesting consistent neural signatures that remain stable across concentration ranges. In contrast, 6:2 FTOH shows significant representation changes across concentrations, making cross-concentration classification more difficult and suggesting that the neural encoding of this compound may be more concentration-dependent. Our findings address several limitations of current PFAS detection technologies, paving the way for employing a brain-based, cyborg sensor with machine learning algorithms for environmental pollution monitoring.

This sensor has been applied to individual PFAS and their concentrations; however, environmental samples often contain complex mixtures of multiple PFAS at varying levels. By incorporating pollutant samples collected from environmental sites, we could evaluate the sensor’s ability in distinguishing PFAS from naturally occurring chemicals. Moreover, by integrating a large, well-characterized computational model, the sensor would be capable of identifying PFAS in contaminated samples, effectively simulating real-world conditions. This approach supports the development of a real-time, adaptive, mobile cyborg sensor designed for field applications, that is highly effective at detecting low-concentration PFAS *in-situ*. Additionally, this technology has the potential to revolutionize chemical sensing by enabling the detection of compounds currently beyond the reach of conventional methods. The locust olfactory system’s vast encoding capacity, precise chemical discrimination, and advanced neural processing make it an exceptionally powerful and adaptable platform for detecting a wide range of chemical substances.

## Acknowledgements

We thank the Michigan State University PFAS Center for suggestions on PFAS and experiments.

## Author contributions

Conceptualization: DS; Formal Analysis: SMS, SJ, AMS, CS; Funding Acquisition: DS; Investigation: SMS; Resources: DS, MB; Writing: SMS, SJ; Review and Editing: DS, AMS.

## Data and Materials Availability

All data, code, and materials will be provided upon request.

## Materials and Methods

### Locust husbandry

Locusts (*Schistocerca americana*) were raised in a crowded colony and kept in an incubator with a 12-hour day and night cycle. The incubator simulated natural temperature fluctuations, maintaining 36.5 ᵒC during the day and 25 ᵒC at night. The locusts were fed a daily mix of fresh grass and wheat germ. Locust eggs were kept in a separate incubator, maintained at a temperature of 37.7 ᵒC.

### Locust surgery

Both sexes of post-fifth instar locusts were used for electrophysiological recordings(*25, 26*). The limbs and wings were removed, and the openings were sealed with Vet bond. The locust was then immobilized on a custom-made surgical platform, secured with electrical tape. Batik wax was used to immobilize the antennae by first constructing a wax pillar on either side of the head, and then securing the antennae into the pillars using tubes. A bowl of batik wax was formed around the head to hold a room-temperature saline solution, ensuring the brain remained moist during excision. A two-part epoxy was used to stabilize the antennae. The head’s exoskeleton was carefully cut to remove the glandular tissue, exposing the brain and gut. The gut was then removed, and a platform was inserted to stabilize and lift the brain. Finally, the ALs were desheathed using a protease treatment.

### Odor vial preparation

For the pure PFAS, the following seven odorants were used: 6:2 FTOH, 8:2 FTOH, 6:2 FTAcr, EtFOSE, MeFOSE, MeFOSA, and PFOSA. These odorants were distributed into individual vials at a concentration of 100 µg (or the equivalent volume in µL, depending on their state of matter), ensuring an equal volume of each odor. An empty vial was used as a control.

For varying concentrations of the three selected PFAS – 6:2 FTOH, 6:2 FTAcr, and PFOS – each odorant was diluted in 10 mL of ddH_2_O at a concentration of 0.01% vol/vol. Serial dilutions were then performed in 10 mL of ddH_2_O to achieve a concentration of 0.0001% vol/vol. A vial containing only 10 mL of ddH_2_O was used as a control. Raoults Law was used to calculate the ppb and ppt of concentrations, using the density of ddH_2_O. Chemical characteristics (i.e., vapor pressure, etc.) needed for Raoults Law were taken from averaging values found in literature(*51–60*).

### Electrophysiology

*In-vivo* extracellular electrophysiology recordings were performed on either sex of post-fifth instar locusts. A commercial 16-channel Neuronexus silicon electrode (A2×2-tet-3mm-150-150-121) was used to extract neural activity from the ALs during odor stimulus presentation. The electrodes were electroplated prior to recordings to achieve impedances between 200 and 300 kΩ. A silver-chloride reference wire was placed in the saline solution within the wax bowl surrounding the locust’s head to ground the experiments. The electrode was inserted approximately 100 µm into the ALs to collect voltage signals. Extracellular recordings were sampled at 20 kHz from 8 channels, comprising two tetrodes. Neural activity was digitized using an Intan pre-amplifier board (C3334 RHD 16-channel head stage), which transmitted data to an Intan recording controller (C3100 RHD USB interface board) (**fig. S1**). The neural activity throughout odor presentation was visualized and stored via the Intan graphical user interface and a LabView data acquisition system, which synchronized both odor delivery and recording windows (**fig. S1B**). For pure PFAS electrophysiology recordings, the same stimulus panel was used, consisting of 7 pure PFAS and a control vial. For concentration experiments, the stimulus panel consisted of 3 PFAS, each diluted to two separate concentrations (0.01% and 0.0001% vol/vol), for a total of 6 odors, plus a control vial. Each odor panel was pseudorandomized before recordings, with 5 trials per odor and a 1-minute interstimulus interval. A total of 13 locusts were used for pure PFAS experiments, resulting in 29 AL recordings. From those recordings, 86 neurons were spike-sorted and used for classification analysis. For the concentration experiments, 8 locusts were used, yielding 18 AL recordings. From these recordings, 69 neurons were spike-sorted. R.M.S. voltage traces were used for classification analysis.

### Odor stimulation

Odor stimulus presentation followed pre-established methodology(*24, 27*). Briefly, a commercial olfactometer (Aurora Scientific, 220A) controlled the odor presentation and airflow (**fig. S1**). A fresh air line delivered 200 sccm of contaminant-free air through a 1/16” diameter PTFE stimulus flow line to the locust antennae, while 200 sccm of air was directed through the dilution flow line to exhaust. The end of the stimulus flow line was position 2 – 3 cm from the antennae’s distal segment. Approximately 5 seconds before odor stimulus, 40% (80 sccm) of the dilution flow was redirected through the odor flow line, upstream of the odorant vials. Both lines merged downstream of the vials, mixing with 120 sccm of clean air from the dilution flow, creating a total of 200 sccm. This allowed a volatile mixture from the stimulus odor to prime the lines up to the final valve, which was previously directed to the exhaust. Once activated, the final valve switched the fresh air flow to exhaust and directed the volatile mixture to the stimulus flow line, delivering the odor to the locust antennae. The odor stimulus lasted 4 seconds before the final valve diverted fresh air to the antennae and exhausted the volatile mixture. One second after stimulus removal, the 80 sccm flowing through the odor flow line was rerouted to prevent potential depletion of the odorant in the vial. A 6” diameter funnel, generating a slight vacuum, was placed directly behind the locust antennae to remove lingering odorants. The odor stimulation protocol was designed to maintain consistent air flow, minimizing potential responses due to changes in air pressure.

### Spike sorting

A high-pass Butterworth filter, set to 300 Hz, was applied to remove low-frequency components. The raw neural data was imported and analyzed using custom-written codes in MATLAB R2024a scripts. The data was then converted into a format readable by IGOR Pro 4 for spike-sorting analysis, following previously described methods to isolate active neurons during stimulus windows(*24, 27*). Spike-sorting was performed with detection thresholds set between 2.5 to 3.5 standard deviations (SDs) from baseline fluctuations. Neurons were considered acceptable if they met the following criteria: < 10% inter-spike intervals (ISIs), spike amplitudes > 5 SDs apart from other sorted neurons, and spike waveform variance < 10%. Additionally, neurons that passed these initial criteria had to exhibit consistent baseline firing for each odor stimulus across five trials, as visualized on raster plots. A total of 86 neurons were isolated for pure PFAS recordings from 13 locust, and 69 neurons were isolated for PFAS concentration recordings using spike sorting.

### RMS transformation

The raw data was imported into MATLAB and processed using a 300 Hz filter. Artifacts caused by electrical interference were identified and removed, resulting in 180 samples centered around each artifact, which were then normalized to the mean voltage value. This process effectively eliminated any electrical interference artifacts from the raw data. For pure PFAS, of the total recorded RMS responses (2,040 Samples: 51 tetrodes x 8 odors x 5 trials), only 29 voltage traces contained artifacts that required removal. No artifacts were found in the PFAS concentration data; therefore, no voltage samples were discarded. The filtered data was then trimmed to the time window of interest before being processed through an R.M.S. filter using previously described methods(*24, 27*). Briefly, a 500-point continuously moving R.M.S. filter, accompanied by a smoothing, 500-point continuously moving averaging filter were applied to the voltage data. Baseline values were calculated by averaging all time bins and trials from the two seconds prior to odor presentation. The data was then normalized by subtracting these baseline values to secure ΔR.M.S. values. These values were binned into 50 ms non-overlapping time bins, and the average of each bin was calculated. For pure PFAS experiments, R.M.S. transformed voltage data from four channels on each tetrode were averaged over 29 positions (51 tetrodes) from 13 locusts (**fig. S2**). A similar procedure was applied to the recorded 18 positions (34 tetrodes) from 8 locust in the PFAS concentration experiments. R.M.S. data were used for analysis in **figures 3 and 4**.

### Dimensionality reduction analysis

Two dimensionality reduction techniques – principal component analysis (PCA) and linear discriminant analysis (LDA) – were used to qualitatively visualize odor-evoked responses in neuronal activity. In PCA, each neuron’s baseline response was calculated by averaging the firing rate during the two seconds before odor presentation. The binned baseline was then subtracted from the spike-sorted neuronal activity. Neuron signals were binned in 50 ms non-overlapping time intervals and averaged across five trials, each with a one-minute interstimulus interval. All spike-sorted neurons were pooled across multiple experiments. For example, in Fig. 2a, the binned responses of all recorded neurons (n = 86) and the number of 50 ms bins between 250 ms and 1000 ms were combined into a matric with neuron number (86) x time bins (t = 15). This matrix represented the spike count of each neuron in each 50 ms time bin. This approach was applied to the entire neuronal time-series population, capturing changes in neural trajectory evolution over time for each odor stimulus. PCA was performed on the time-series data from all seven pure PFAS and the control, maximizing the variance within the data set. High-dimensional time bin vectors were projected onto the principal component axes to visualize the directions of maximum variance. The three-dimensional vectors with the highest eigenvalues were used to plot and connect the data points in adjacent time bins, illustrating neural trajectory evolution over time (**Fig. 2A**). The trajectories converge at the origin to explore the dynamics of odor-specific responses and trajectory separation. PCA analysis using baseline-subtracted, R.M.S. transformed population data was also applied to the pure PFAS data (**fig. S2A**). A similar PCA analysis was applied to the PFAS concentration data using baseline-subtracted, R.M.S. transformed population data (**Figs. 3D and 4D, fig. S3C**). For LDA, analogous neural population data matrices were used for the evaluation of pure PFAS (**Fig. 2B**), and R.M.S. transformed population data was further used for both pure PFAS and PFAS concentration data (**Figs. 3C and 4C, figs. S2B and S3B**). LDA minimizes within-stimulus variation while maximizing the separation between stimuli, highlighting the differentiation in spatio-temporal space for each odor. The time bins were plotted as points in the LDA space, revealing distinct neural clustering for stimulus response. All dimensionality reduction analyses were performed using custom-written MATLAB R2024a scripts.

### LOTO confusion matrix

To quantitatively classify individual PFAS concentrations, we employed a leave-one-trial-out (LOTO) confusion matrix. This method evaluates the accuracy of PFAS classification within our model, demonstrating how effectively locust neural responses differentiate between PFAS, various concentrations of the same chemical, and an odor control. After spike sorting, neuronal population activity is divided into 50 ms time bins, creating a three-dimensional matrix (e.g., 86 neurons x 5 trials x 110-time bins). The number of bins depends on the time window of interest; for this data, the time window was 0.25 – 5.75 s (5,500 ms) after odor presentation. This time window encompasses both the ON and OFF responses, capturing the entire neuronal odor-evoked response. The spiking events in each time bin are counted for all neurons in the population. This process is repeated for each neuron across five trials. Four of the five trials are averaged to create a training template (predicted label), while the fifth trial (the left-out trial) serves as a testing template (true label). The number of predicted and true labels corresponds to the number of odors (e.g., for 8 odors, there will be 8 predicted and 8 true labels). These templates are plotted in high-dimensional space, and each testing template is assigned to a training template based on the smallest Euclidean distance. A confusion matrix is then generated to depict classification accuracy of the assignments. Using a “winner-takes-all” approach, or the mode of time bin assignments, the testing trials are matched to their respective training templates. This results in a trial-wise LOTO confusion matrix that indicates the accuracy of odor classification. A fully diagonal matrix indicates 100% classification accuracy. A similar analysis was performed for R.M.S. transformed pure and concentration PFAS data (**Figs. 3E and 4E, figs. S2C and S3D**). In this case, the three-dimensional matrix was based on the number of tetrodes instead of neurons (e.g., 34 tetrodes x 5 trials x 110-time bins). All quantitative classification analyses were carried out using custom written MATLAB R2024a scripts.

### Sensitivity and Specificity Tables

Values used to calculate sensitivity and specificity were obtained from the LOTO confusion matrices. The columns of the matrix were divided between true positives (TP) and false negatives (FN), while the rows were split between false positives (FP) and TP. Any data not in the odor column or row of interest was considered a true negative (TN). A TP was identified as the diagonal classification accuracy, where the true label was correctly assigned to the predicted label.

Sensitivity was calculated as the ratio of TP to the sum of TP and FN, representing all positive conditions.

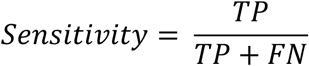

Specificity was calculated as the ratio of TN to the sum of TN and FP, highlighting all negative conditions.

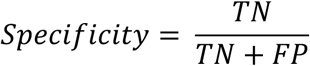

For calculations of groups (i.e., PFAS vs Air, PFAS families, PFAS vs ddH_2_O), all odors of interest values were combined. For the FTOH family, TP values were defined as any classification that was accurately assigned to either 6:2 FTOH or 8:2 FTOH, including the misclassification of one as the other (**Figs. 2C and 2D**). False positives (FP) were any odors outside the FTOH family assigned to either FTOH compound. True negative (TN) were considered as any odor assignments not in the FTOH family. A similar process was applied to other PFAS families analyzed (**Fig. 2D and fig. S2D**). For the PFAS vs Controls groupings, any assignment of a PFAS odor to a PFAS odor were considered a TP. Assigning a PFAS odor to a control was a FN, and assigning a control to a PFAS odor was a FP. TN were defined as an accurate assignment of the control odor to itself. Analogous sensitivity and specificity tables were created for PFAS concentrations (**Figs. 3F and 4F, fig. S3E**).

### Machine Learning Model

We implemented a machine learning algorithm for PFAS classification and automated identification of unknown samples. Neural activity (R.M.S. data) from locust AL recordings during PFAS exposure were preprocessed using standardized z-scoring across all training odors and trials. We evaluated two classification paradigms: (a) classification of odor identity only, ignoring concentration and (b) generalization across concentrations, training on high and testing on low concentrations. For (a), both low and high concentrations were included in the training data.

We implemented a CNN-LSTM (Convolutional Neural Network – Long Short-Term Memory) hybrid model employing two sequential, one-dimensional convolutional layers with increasing feature dimensions to extract hierarchical spatial patterns from neural responses. This was followed by a single-layer LSTM for temporal feature extraction and modeling of neural dynamics over time. The architecture incorporated a two-layer classification head using ReLU activations and dropout regularization to prevent overfitting. Classification performance was evaluated using leave-one-trial-out cross-validation for the first classification experiment. For generalization across concentrations, we used neural responses from high concentration compounds for training and tested on responses from the same compounds at lower concentrations. The results were averaged across 50 random initializations to provide statistical confidence in reported accuracies.

The data was z-scored across all odors and trials for all the experiments. For the generalization experiment, the data was normalized based on the maximum channel activation for each trial and odor in each time bin.

