## Supplemental Data for "Detection of multiple per- and polyfluoroalkyl substances (PFAS) using a biological brain-based gas sensor"

**Affiliations:**
[1]Department of Biomedical Engineering, Michigan State University, East Lansing, MI

[2]Institute for Quantitative Health Sciences and Engineering, Michigan State University, East Lansing, MI

[3]Department of Electrical and Computer Engineering, University of California San Diego, La Jolla, CA

[4]Department of Medicine, University of California San Diego, La Jolla, CA

[5]Neuroscience Program, Michigan State University East Lansing, MI

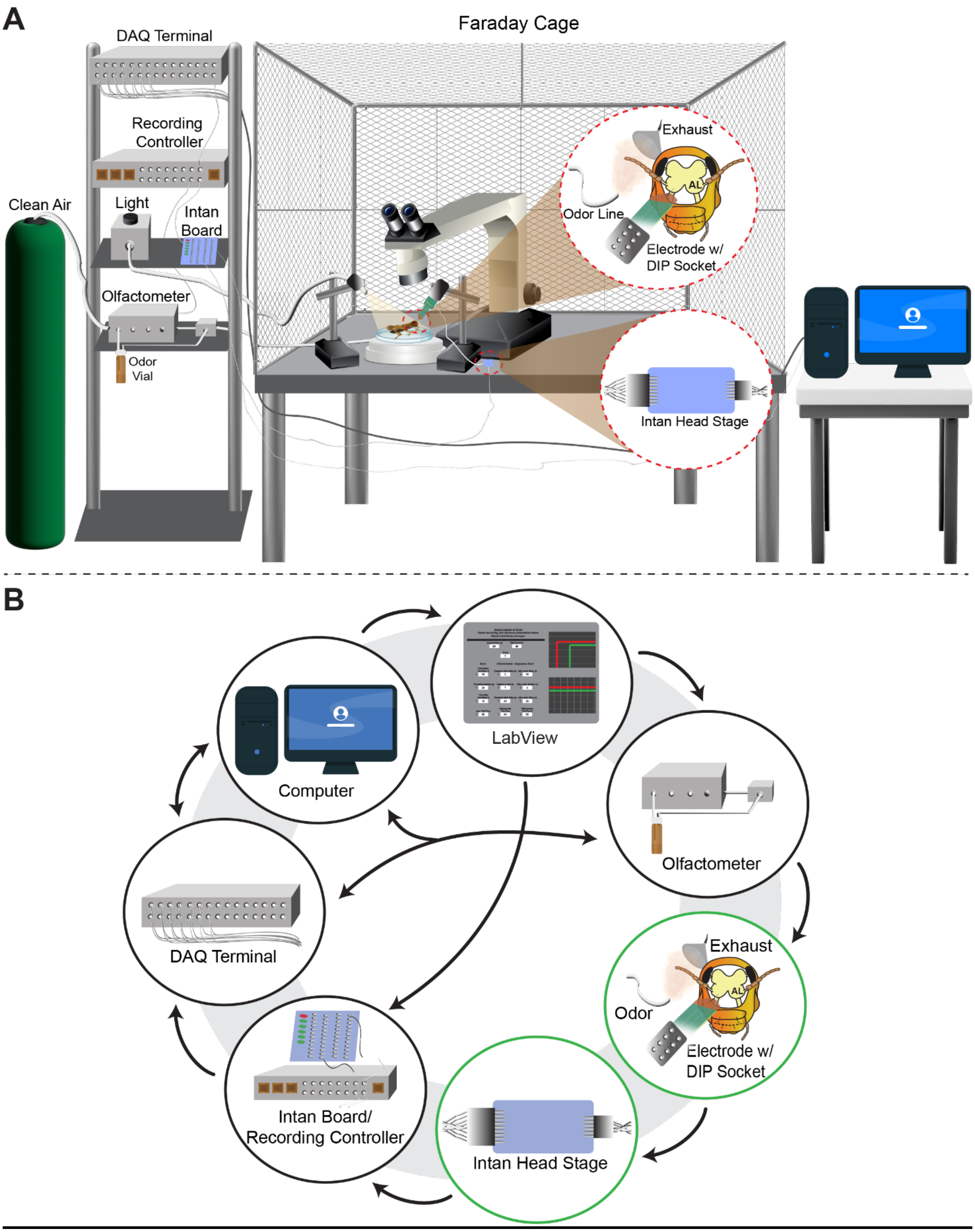


**fig. S1. Overview of electrophysiology design and signaling pathways. (A)** Clean air is mixed with odor laden air using the olfactometer before delivering to the locust antennae via an odor line.

Odor presentation timing is controlled through the olfactometer, recording controller, and Intan board. Information is relayed through the DAQ terminal to the computer. The faraday cage houses the insect with electrode implanted in the AL and the Intan head stage to isolate the recordings, limiting electrical interference. Neural activity is visualized and stored on the computer. **(B)** Diagram depicts device signaling pathways with directional arrows indicating communication from one device to the next. Bi-directional arrows indicate communication both ways. Green circles designate devices found within the faraday cage.

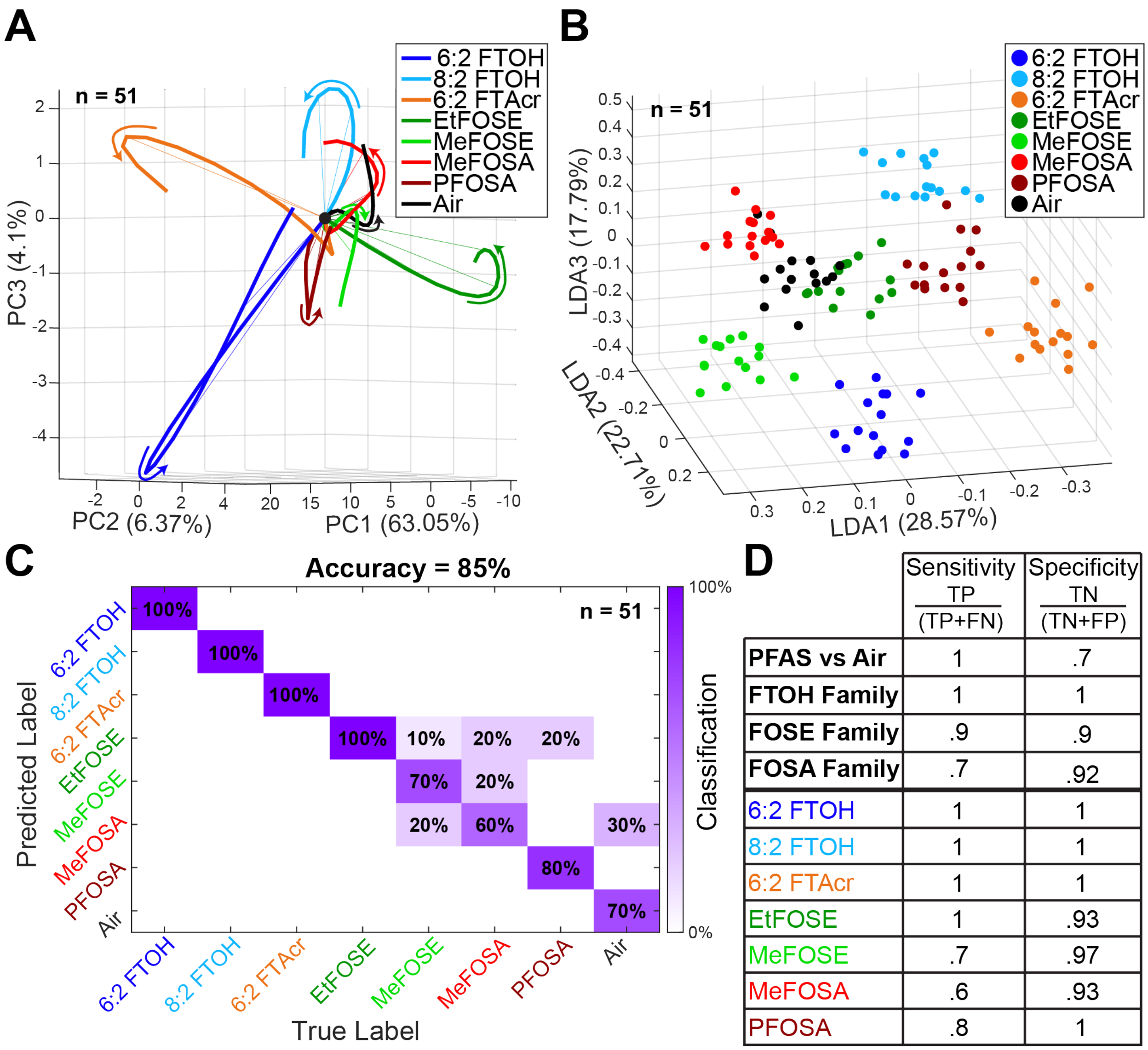


| | Sensitivity $\frac{TP}{(TP+FN)}$ | Specificity $\frac{TN}{(TN+FP)}$ |
|---|---|---|
| **PFAS vs Air** | 1 | .7 |
| **FTOH Family** | 1 | 1 |
| **FOSE Family** | .9 | .9 |
| **FOSA Family** | .7 | .92 |
| 6:2 FTOH | 1 | 1 |
| 8:2 FTOH | 1 | 1 |
| 6:2 FTAcr | 1 | 1 |
| EtFOSE | 1 | .93 |
| MeFOSE | .7 | .97 |
| MeFOSA | .6 | .93 |
| PFOSA | .8 | 1 |

**fig. S2. Classification of pure PFAS using R.M.S. analysis. (A)** Odor-evoked R.M.S. trajectories for all seven pure PFAS are visualized using PCA for dimensionality reduction. These trajectories represent neural activity from 0.25 – 1.0 seconds following odor exposure. The colored arrows depict the progression of trajectories, aligned at 0.25 seconds after odor onset (black dot). R.M.S captures the real-time evolution of odor responses through the sensor. A total of 51 tetrodes were analyzed. **(B)** LDA, a 3-class supervised dimensionality reduction technique, maps spatiotemporal R.M.S. odor responses between 0.25 – 1.0 seconds post-exposure. Clustering reveals distinct temporal patterns for each odor stimulus (n = 51). **(C)** A high-dimensional LOTO confusion matrix classifies R.M.S. responses to the seven pure PFAS and the control. Classification of true versus predicated labels was based on the smallest Euclidean distance within the 0.5 – 6.0 second window after stimulus onset. **(D)** Sensitivity and specificity table depicting real-time compound detection. Values were generated from the LOTO matrix (Materials and Methods).

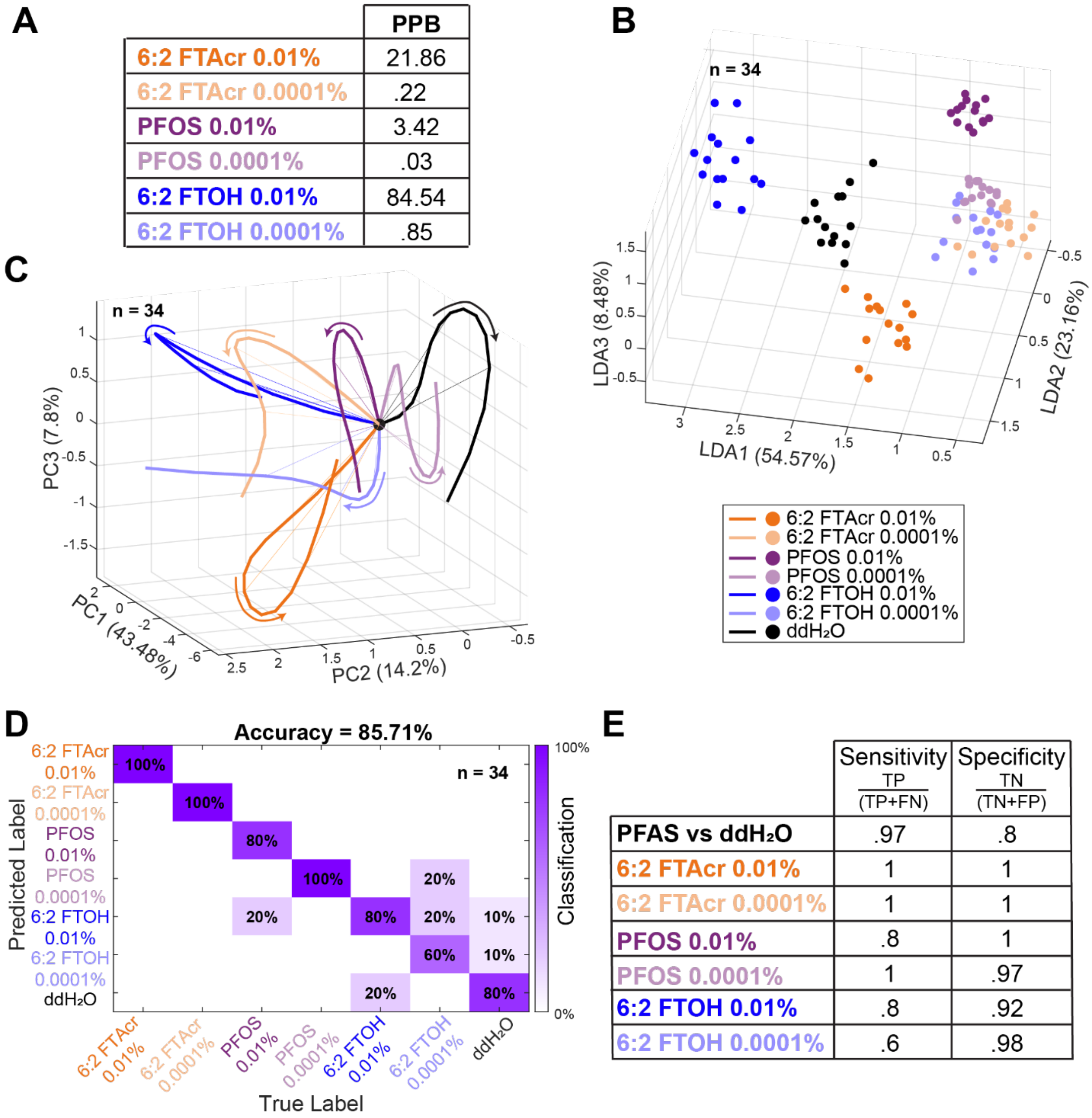


| | PPB |
|---|---|
| 6:2 FTAcr 0.01% | 21.86 |
| 6:2 FTAcr 0.0001% | .22 |
| PFOS 0.01% | 3.42 |
| PFOS 0.0001% | .03 |
| 6:2 FTOH 0.01% | 84.54 |
| 6:2 FTOH 0.0001% | .85 |

| | Sensitivity TP/(TP+FN) | Specificity TN/(TN+FP) |
|---|---|---|
| PFAS vs ddH₂O | .97 | .8 |
| 6:2 FTAcr 0.01% | 1 | 1 |
| 6:2 FTAcr 0.0001% | 1 | 1 |
| PFOS 0.01% | .8 | 1 |
| PFOS 0.0001% | 1 | .97 |
| 6:2 FTOH 0.01% | .8 | .92 |
| 6:2 FTOH 0.0001% | .6 | .98 |

**fig. S3. Environmental PFAS concentrations show high classification accuracy using R.M.S. (A)** Ppb calculations for three PFAS at two concentrations (6 odors total) were made using Raoult's law and published chemical characteristics (Materials and Methods). The lowest concentration (0.0001%) represents ppt ranges. **(B)** A 3-class LDA, a supervised dimensionality reduction technique based on R.M.S., effectively separates odors and the control into distinct clusters. The stimulus window analyzed was 0.25 – 1.0 seconds post-odor onset (n = 34 tetrodes). **(C)** PCA dimensionality reduction highlights the separation of R.M.S neural trajectory evolution from 0.25 – 1.0 seconds after stimulus onset. Colored arrows illustrate the trajectory evolution, aligned at 0.25 seconds after odor presentation (black dot). Trajectories for different concentrations of a single odor follow similar responses (n = 34 tetrodes). **(D)** A LOTO confusion matrix classifies R.M.S. responses of the PFAS concentrations, along with a control, achieving 86% accuracy. Classification assignments were based on the smallest Euclidean distance within the 0.25 – 5.75 second window after stimulus onset. **(E)** A sensitivity and specificity table demonstrates real-time PFAS concentration detection. Values were generated from the LOTO matrix (Materials and Methods).
